# The Effect of Minnelide against SARS-CoV-2 in a Murine Model

**DOI:** 10.1101/2021.05.05.442875

**Authors:** Marley C. Caballero Van Dyke, Heather L. Mead, Mitchell Bryant, Klaire L. Laux, Daniel R. Kollath, Vanessa K. Coyne, Karis J. Miller, Nathan E. Stone, Sierra A. Jaramillo, Paul Keim, Clara Milikowski, Vineet K. Gupta, Selwyn M. Vickers, Ashok K. Saluja, Mohana R. Velagapudi, Bridget M. Barker

**Author notes:** **Address Correspondence to** Mohana Velagapudi, M.D.

## Abstract

Severe acute respiratory syndrome coronavirus 2, SARS-CoV-2, is the causative agent of coronavirus disease 2019, COVID-19, and the current COVID-19 pandemic. Even as more vaccine candidates are released, more treatment options are critically needed. Here, we investigated the use of Minnelide, a water soluble pro-drug with anti-inflammatory properties, for the treatment of COVID-19. To do this, k18-hACE2 mice were infected with SARS-CoV-2 or given PBS control intranasally. The next day mice were either treated daily with low dose (0.0025mg/day) or high dose Minnelide (0.005mg/day), or given vehicle control intraperitoneal. Mice were weighed daily, and sacrificed at day 6 and 10 post-infection to analyze viral burden, cytokine response, and pathology. We observed a reduction in viral load in the lungs of Minnelide-treated mice infected with SARS-CoV-2 at day 10 post-infection compared to day 6 post-infection. All SARS-CoV-2 infected non-treated mice were moribund six days post-infection while treatment with Minnelide extended survival for both low (60% survival) and high (100% survival) dose treated mice ten days post-infection. Interestingly, cytokine analysis demonstrated a significant reduction in IL-6 (lung and heart) and D-dimer (serum) in high dose treated SARS-CoV-2 infected mice compared to mice infected with SARS-CoV-2 alone at day 6 post-infection. Additionally, histology analysis revealed that Minnelide treatment significantly improved lung pathology ten days post-infection with SARS-CoV-2 with all the mice exhibiting normal lung tissue with thin alveolar septa and no inflammatory cells. Overall, our study exhibits potential for the use of Minnelide to improve survival in COVID-19 patients.

## 1 Introduction

Coronavirus disease 2019 (COVID-19) is an infectious respiratory disease caused by a novel severe acute respiratory syndrome coronavirus 2 (SARS-CoV-2) [1]. This novel viral disease was first recognized in December 2019 in Wuhan City, China and has since spread rapidly throughout the world, and subsequently causing the worst global pandemic of recent times with more than 140 million confirmed cases and more than 3 million deaths worldwide [2]. Patients with COVID-19 most commonly report fever, chills, fatigue, cough, shortness of breath, and difficulty in breathing which may appear 2-14 days after exposure to the virus. In some cases, patients also have increased sputum production, anosmia, dyspnea, ageusia, headache, hemoptysis, diarrhea, and myalgia [3, 4]. Interestingly, roughly 20% percent of patients did not show any symptoms but still have capacity to transmit disease to other healthy individuals making this disease difficult to control [5]. In general, adults with SARS-CoV-2 infection can be grouped into the following severity of illness categories [6]. *Asymptomatic or Presymptomatic Infection*: Individuals who test positive for SARS-CoV-2 using a virologic test (i.e., a nucleic acid amplification test or an antigen test) but who have no symptoms that are consistent with COVID-19. *Mild Illness*: Individuals who have any of the various signs and symptoms of COVID-19 (e.g., fever, cough, sore throat, malaise, headache, muscle pain, nausea, vomiting, diarrhea, loss of taste and smell) but who do not have shortness of breath, dyspnea, or abnormal chest imaging. *Moderate Illness*: Individuals who show evidence of lower respiratory disease during clinical assessment or imaging and who have saturation of oxygen (SpO2) ≥94% on room air at sea level. *Severe Illness*: Individuals who have SpO2 <94% on room air at sea level, a ratio of arterial partial pressure of oxygen to fraction of inspired oxygen (PaO2/FiO2) <300 mm Hg, respiratory frequency >30 breaths/min, or lung infiltrates >50%. *Critical Illness*: Individuals who have respiratory failure, septic shock, and/or multiple organ dysfunction. However, the criteria for each category may overlap or vary across clinical guidelines and clinical trials, and a patient’s clinical status may change over time. Mortality among these patients mainly occurs through the development of viral pneumonia-induced acute respiratory distress syndrome (ARDS).

In ARDS, SARS-CoV-2 infection triggers a dysfunctional immune response causing a cytokine storm which causes most of the damage to the lung. COVID-19 patients with ARDS have higher level of proinflammatory cytokine like interleukin (IL)-1α, IL-1β, IL-1 receptor antagonist protein (IL1RA), IL-2, IL-4, IL-6, granulocyte-macrophage colony-stimulating factor (GM-CSF), and interferon (IFN)-gamma in blood compared to healthy controls [4, 7]. Further analysis of Bronchoalveolar lavage fluid (BALF) in patients with severe COVID-19 also showed higher levels of IL-6, IL-8, and IL-1β [8]. Single-cell RNA sequencing analysis of BALF showed enrichment of proinflammatory monocyte-derived macrophages in patients with severe COVID-19 [8] resulting in macrophage-activation syndrome (MAS), a life-threatening clinical manifestation observed in autoimmune diseases and several viral infections like influenza [9]. Macrophage activation often leads to impairment of cytotoxic activity of CD8^+^ T cells and NK cells thereby preventing clearance of infected host cells [10]. In addition, SARS-CoV-2 infection also activates TNF and IFN signaling thereby causing T-cell apoptosis, further causing hyperinflammation and virus replication, and subsequently, causing damage to lung epithelial and endothelial cells resulting in ARDS.

SARS-CoV-2 is a positive-sense, single-stranded RNA viruses which belong to the Betacoronavirus genus [11]. These viruses have spike (S)-glycoprotein on their surface, which through the receptor-binding domain (RBD) facilitates the entry of SARS-CoV-2 into host cells by binding to the human angiotensin-converting enzyme 2 (ACE2) receptor [12]. ACE2 is predominantly expressed in the epithelium of the human lung, small intestine, and oral mucosa [12, 13] and therefore provide route of infection. Once inside the cell, the viral genome is transcribed by the viral RNA-dependent RNA polymerase and subsequently translated using host ribosomes to synthesize viral proteins. Finally, mature virions are assembled and packaged in the cytoplasm of the infected cell and subsequently released through exocytosis. ACE2 expression often determines the susceptibility for COVID-19 infection. For example, children express very low level of ACE2 which makes them less susceptible to SARS-CoV-2 infection compared to adults [14]. Notably, structural differences between mouse ACE2 and the human counterpart makes it unsuitable for the spike protein of SARS-CoV-2 and therefore it does not facilitate the entry of the virus into the cells. As a result, wild-type laboratory mice are not suitable for SARS-CoV-2 infection studies. However, new studies demonstrate the ability of variants of concern (VOCs) to infect common laboratory mice [15]. To study SARS-CoV-2 pathogenesis and potential treatment options, the ideal model is the K18-hACE2-transgenic mice because SARS-CoV-2 infection in these mice replicates human pulmonary disease similar to COVID-19. This is due to human-like ACE-2 expression driven by the epithelial cell cytokeratin-18 (K18) promoter, which expresses ACE-2 in epithelial cells [16]. This mouse model was initially developed by Tseng, C. *et al* [17] for the study of SARS-CoV pathogenesis. SARS-CoV-2 also uses ACE-2 for the entry of virus into the host cell, and mimics respiratory disease similar to severe COVID-19 in humans [18].

Minnelide is a water soluble pro drug of an anti-inflammatory active diterpene triepoxide compound triptolide which was purified from a Chines herb Tripterygium wilfordii Hook F [19]. Historically, triptolide has been used for the therapy of arthritis in traditional Chinese medicine for more than two thousand years. Triptolide has been documented to decrease inflammation by decreasing NF-kB activity [20–22] leading to a decrease in production of inflammatory cytokines like IL-6 [23] and IL-1β [24]. Triptolide, the active component of this drug is also reported to inhibit acute lung injury in murine models [25]. Additionally, unpublished preclinical studies from our laboratory show that in a mouse model of pulmonary fibrosis, Minnelide was highly effective in decreasing fibrosis and showed improvement in various lung functions. In a mouse model of pancreatic cancer, Minnelide showed a decrease in IL-1β as well as NF-kB activity [20, 26]. Anti-inflammatory and anti-fibrotic properties of Minnelide not only makes it an ideal candidate for early therapeutic intervention against COVID-19 patients, who develop ARDS because of hyperactive immune response and cytokine storm, but also makes it suitable for depleting fibrosis caused by COVID-19 associated lung injury. In the current study, we show that Minnelide treatment significantly improved survival of K18-hACE2 mice infected with SARS-CoV-2. Further analysis showed significant reduction in SARS-CoV-2 viral burden in lungs of Minnelide treated mice. Our results also show that Minnelide treatment elicited an anti-inflammatory response against SARS-CoV-2 resulting in improved lung histology and thereby highlighting its therapeutic potential in COVID-19 patients suffering from ARDS.

## 2 Results

### 2.1 SARS-CoV-2 Viral Burden in Mice Treated with Minnelide

To understand whether treatment with Minnelide would demonstrate therapeutic benefits against COVID-19, we infected K18-hACE2 mice with 2000 PFU of SARS-CoV-2 or PBS control via intranasal infection. The next day mice were treated daily with either low dose (0.0025mg/day) or high dose (0.005mg/day) Minnelide, or vehicle control, PBS via intraperionteal injection (IP) (Figure 1).

**Figure 1.**
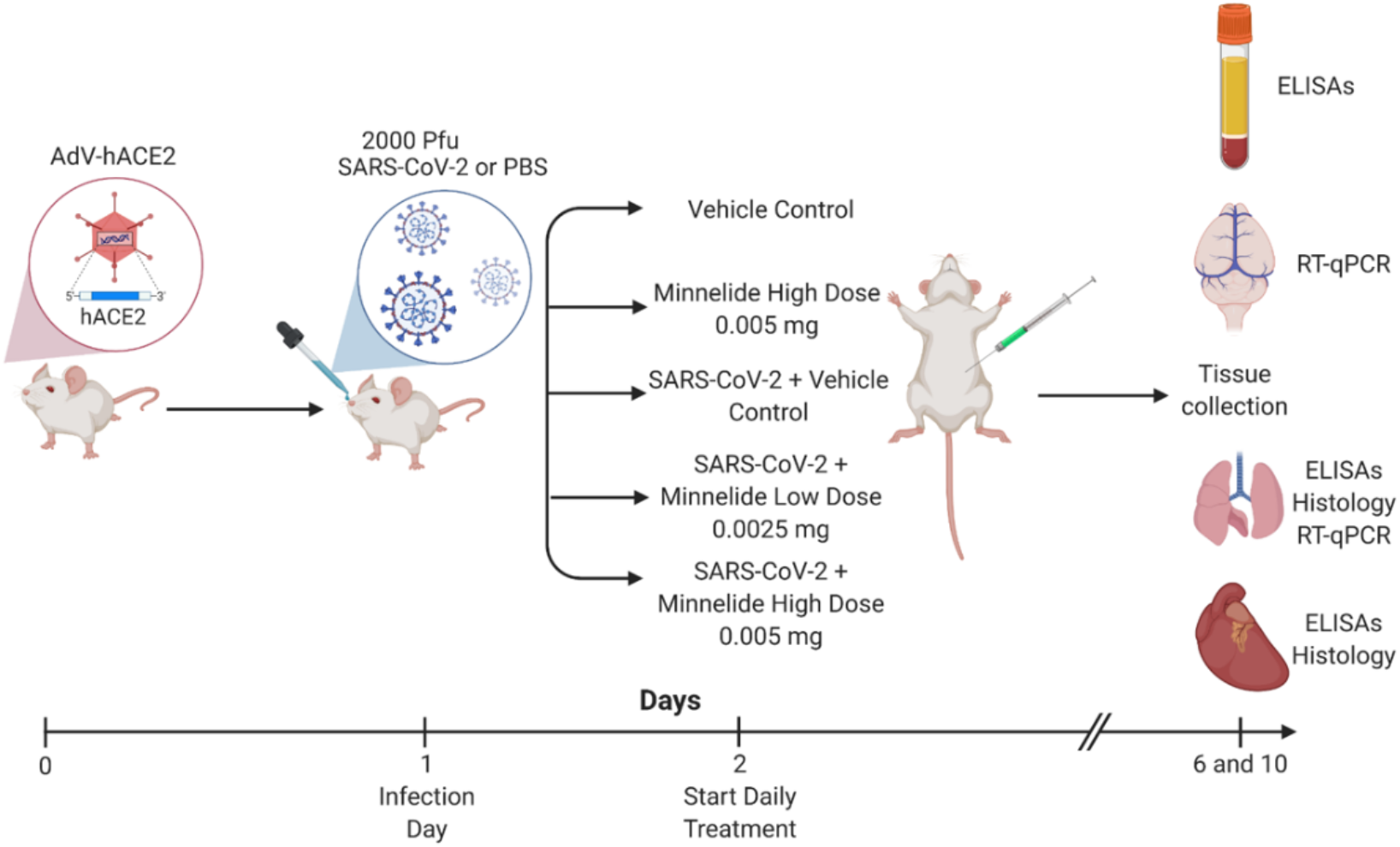
Experimental Schematic of Infection and Treatment of K18-hACE2 Mice. K18-hACE2 were infected with 2000 PFU on day 1 or given PBS intranasally. Daily treatments of Minnelide started on day 2 via intraperitoneal (IP) injections with either vehicle control (PBS), Minnelide low dose (0.0025mg/day or high dose (0.005mg/day). On days 6 and 10, tissues were extracted for ELISAs, RT-qPCR, or Histology.

Viral burden was measured in the lung and brain at day 6 and 10 post-infection using RT-qPCR and compared among groups (Figure 2; Table S1 and S2). In the brain, the highest levels of viral burden were detected in SARS-CoV-2 non-treated mice and low dose treated mice at day 6 post-infection (Figure 2; Table S1). Mice infected with SARS-CoV-2 and treated with high dose of Minnelide still exhibited a high viral load at day 6 post-infection (Figure 2; Table S1). For day 10 post-infection, the brain did exhibit less viral burden in both the low dose and high dose Minnelide treated groups compared to day 6 post-infection, although not significant (Figure 2; Table S1). In the lung, there was high viral load detected at day 6 post-infection across all groups, SARS-CoV-2 infected non-treated mice, low dose, and high dose Minnelide treated mice infected with SARS-CoV-2 (Figure 2; Table S2). At day 10 post-infection, there was a lower level of viral burden in both the low and high dose treated mice infected with SARS-CoV-2, although not significant (Figure 2; Table S2). Together, these results indicate that Minnelide treatment results in a measurable, but not statistically significant reduction in viral burden.

**Figure 2.**
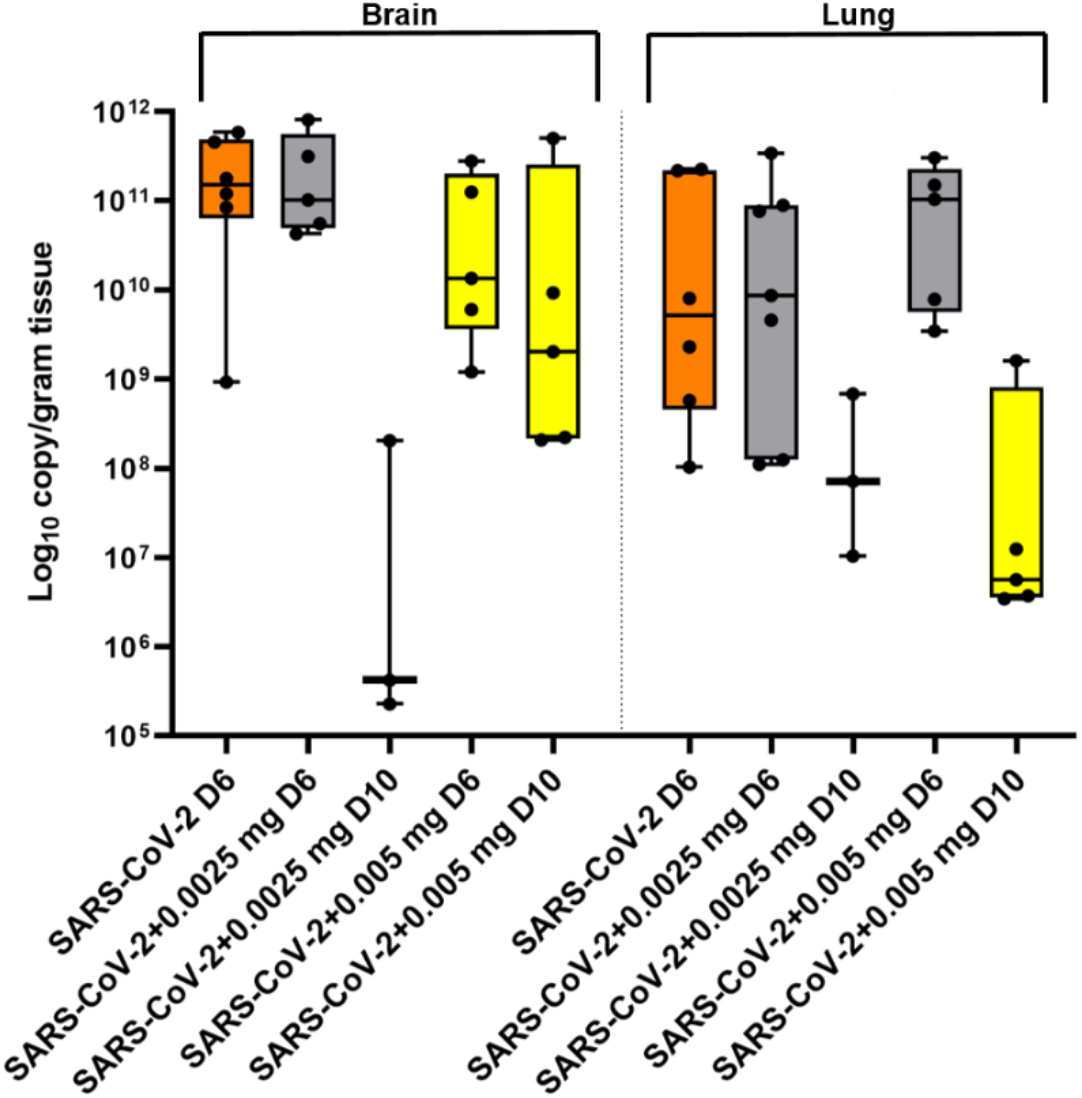
Viral burden of SARS-CoV-2 Infected Mice Treated with Minnelide. K18-hACE2 mice received an intranasal infection with 2000 PFU of SARS-CoV-2 and were treated twice daily via oral gavage with either low dose (0.0025mg) or high dose (0.005mg) of Minnelide. At days 6 and 10 post infection, brain and lung were excised, homogenized, and RNA was extracted. RNA data is cumulative of one experiment using 3-6 mice per group per experiment.

### 2.2 Treatment with Minnelide Increases Survival and Elicits an Anti-Inflammatory Response against SARS-CoV-2

To further understand the therapeutic benefits of Minnelide treatment, we observed mice until 10 days post-infection weighing them daily. SARS-CoV-2 infected non-treated mice demonstrated 100% mortality at day 6 post-infection (Figure 3A). Mice infected with SARS-CoV-2 and treated with low dose Minnelide demonstrated 60% survival by day 10 post-infection. Interestingly, all mice infected with SARS-CoV-2 and treated with high dose Minnelide were alive at day 10 post-infection (100% survival) (Figure 3A). Additionally, we also observed 100% survival rates in the uninfected, vehicle control and Minnelide high dose only treated mice at both time points. This data suggests therapeutic potential of Minnelide for COVID-19 patients.

**Figure 3.**
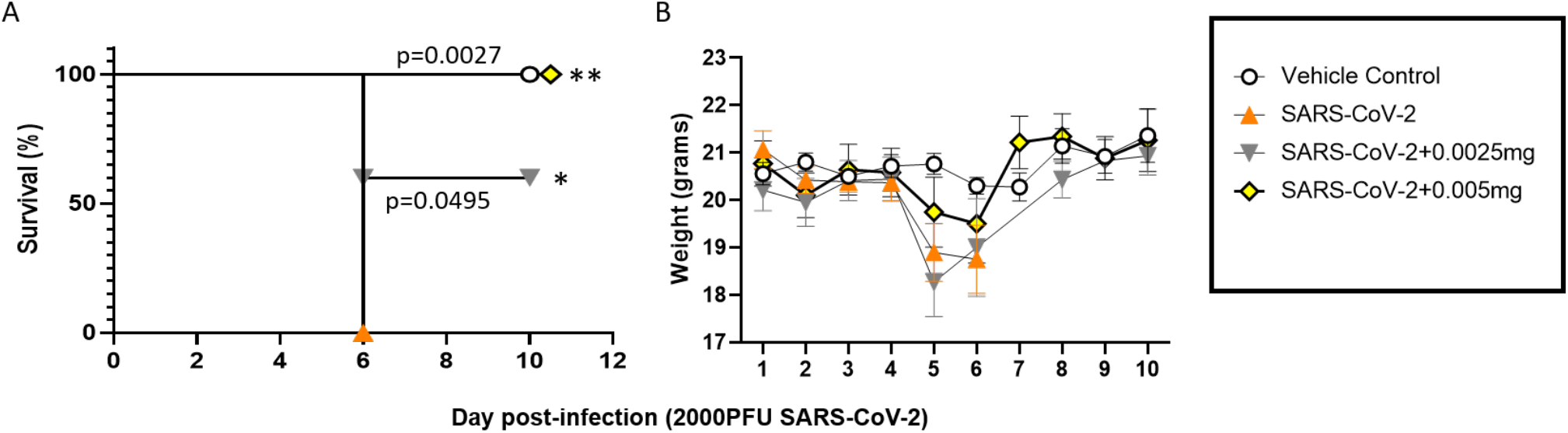
Treatment of Minnelide against SARS-CoV-2 increased protection in mice. K18-hACE2 mice infected with 2000 PFU SARS-CoV-2 and treated with Minnelide were observed for 10 days post-infection for survival analysis (A) and weighed daily (B). For (A), n=5 and (B) n= 5-10 mice per group. Minnelide only group is not presented since this group was female mice and demonstrated much less weight overall, which did not change overtime, compared to male mice. Data are presented as mean values ± SEM (B). Significance was tested by Log-rank (Mantel-Cox) test (A) or one-way ANOVA (B). Survival significance is shown against SARS-CoV-2 non-treated group (A). Data is from one experiment.

Additionally, weights of mice were monitored daily. All non-infected control mice remained within +/- 1g of their starting weight during the course of the experiment (Figure 3B). In contrast, all mice infected with SARS-CoV-2 not treated with Minnelide experienced a significant weight loss between day 4 and day 6 post-infection with no mice surviving past this time point; these mice were moribund (defined as a weight drop of >20%) and euthanized on day six (Figure 3B). The infected mice that received Minnelide therapy experienced less weight loss than infected mice without therapy. The increase in drug dosage, 0.005mg/day treated mice, demonstrated less weight loss compared to low dose, 0.0025mg/day treated mice, although no statistical significance was found (Figure 3B). All mice demonstrated weight loss between days 4-6 post-infection. Interestingly, mice that received treatment recovered their weight after day 6 post-infection. This is clearly demonstrated by the high dose treated mice, because none succumbed to the infection in that group (Figure 3B). All mice infected with SARS-CoV-2 and without Minnelide treatment were euthanized on day 6 post-infection due to extreme weight loss.

Next, we evaluated select cytokines (IL-6, D-dimer, and Fibrinogen) in the lung, heart, and serum in mice infected with SARS-CoV-2 and treated with Minnelide on days 6 and 10 post-infection. For IL-6 in the lungs and heart, we observed a significant reduction in IL-6 production in mice infected with SARS-CoV-2 treated with high dose Minnelide at day 6 post-infection compared to SARS-CoV-2 alone (Figure 4A). We observed no significant differences in IL-6 in the serum across all groups (Figure 4G). For D-dimer, there was a significant reduction in the lungs at day 10 post-infection in both low and high dose treated mice infected with SARS-CoV-2 compared to vehicle control mice (Figure 4B). At day 6 post-infection, we observed a significant reduction in D-dimer in the serum of mice treated with low and high dose Minnelide compared to mice infected with SARS-CoV-2 alone and not treated (Figure 4H). We observed no significant differences in D-dimer production in the heart across groups (Figure 4E). For Fibrinogen, the data show significant reduction at day 10 post-infection in mice treated with high dose Minnelide compared to low dose Minnelide infected with SARS-CoV-2 (Figure 4C). There were no significant differences for Fibrinogen production in the heart or serum across groups (Figure 4F and 4I).

**Figure 4.**
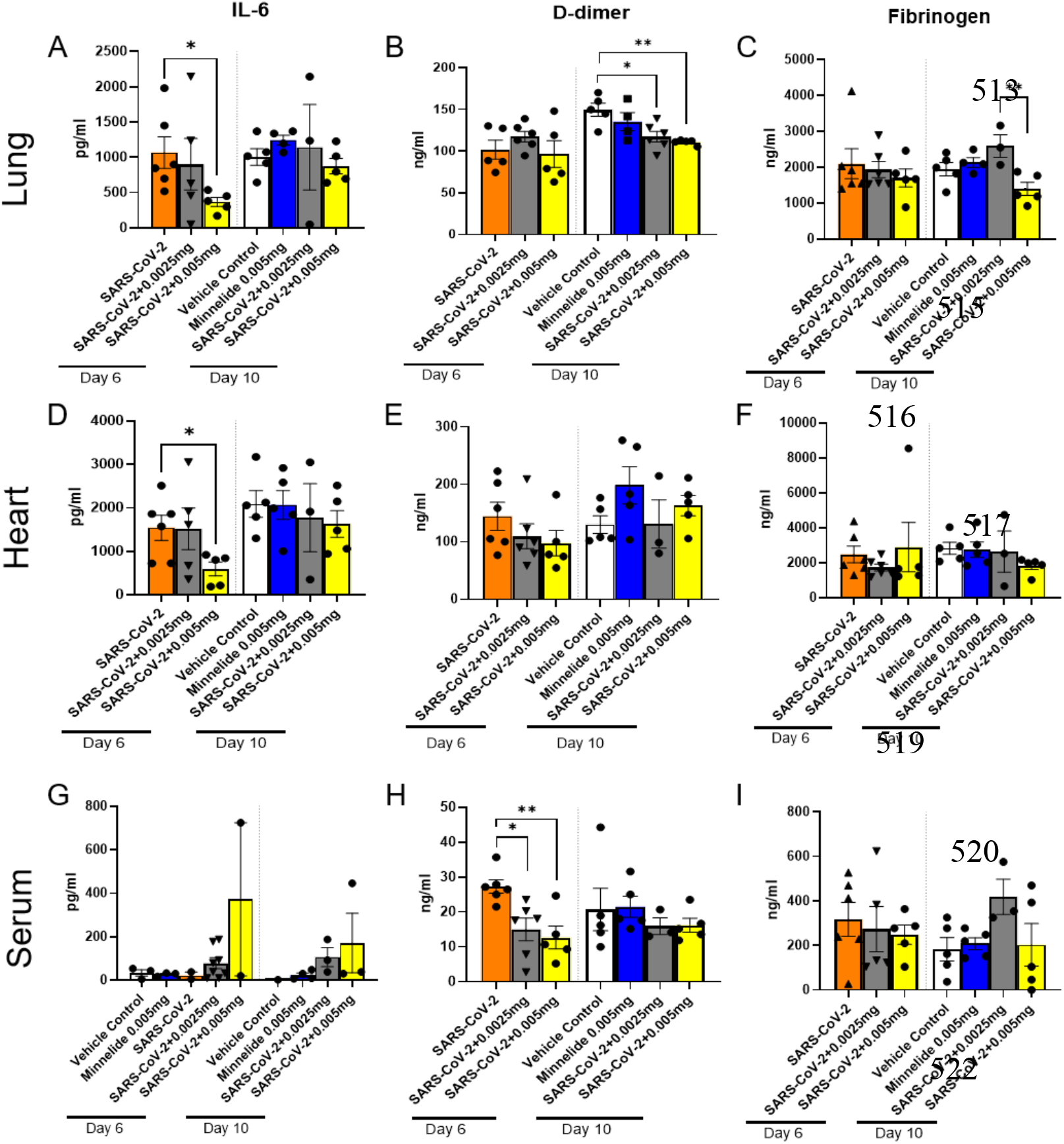
Pulmonary Cytokine Milieu in SARS-CoV-2 Infected Mice Treated with Minnelide. K18-hACE2 mice received an intranasal infection with 2000 PFU of SARS-CoV-2 and were treated daily with either low dose (0.0025mg) or high dose (0.005mg) of Minnelide. At days 6 and 10 post infection, lung (A-C) and heart (B-F) were excised, homogenized, and supernatant collected along with serum (G-I) for cytokine analysis. Cytokine data is cumulative of one experiment using 5-6 mice per group per time point. Significance was tested by one-way ANOVA. * *p* < 0.05 and \*\**p* < 0.001.

### 2.3 Histology

Lastly, we sought to determine the effect of Minnelide treatment on the pathology of mice infected with SARS-CoV-2. Figure 5A is representative of the five mice treated with saline only that were all sacrificed on day 10 and shows normal lung tissue with thin alveolar septa without increase in inflammatory cells. Figure 5B is representative of the five mice that were treated with high dose Minnelide (0.005mg) alone that were all sacrificed on day 10 with similar findings to the saline only group. Figure 5C is representative of the six mice infected with virus without treatment that were sacrificed on day 6. Three of the mice from the virus only group (50%) showed focal widening of alveolar septa with mild increase in small mononuclear cells (lymphocytes) and increased capillaries (Figure 5C). Figures 5D-5I are representative of the group of nine mice (eight mice lungs reviewed) five of which were sacrificed on day 6 (Figures 5D-5F) and four of which were sacrificed on day 10 (Figures 5G-5I) all of which were infected with virus and treated with low dose Minnelide (0.0025mg). On day 6, lungs showed alveolar findings that were similar to the control group with normal findings with the exception of one mouse that showed platelet aggregates in blood vessels (Figures 5E and 5F). Of the three mice that were sacrificed on day 10, all (100%) showed perivascular and peribronchiolar infiltration by large reactive blast-like cells (Figures 5G-5I). Figures 5J-5K are representative of the ten mice infected with virus and treated with high dose Minnelide of which five were sacrificed on day 6 and five were sacrificed on day 10. Alveolar septa were normal in all cases and only one case showed few early fibrin thrombi (Figure 5K). Of those mice sacrificed on day 10, all (100%) were without changes.

**Figure 5.**
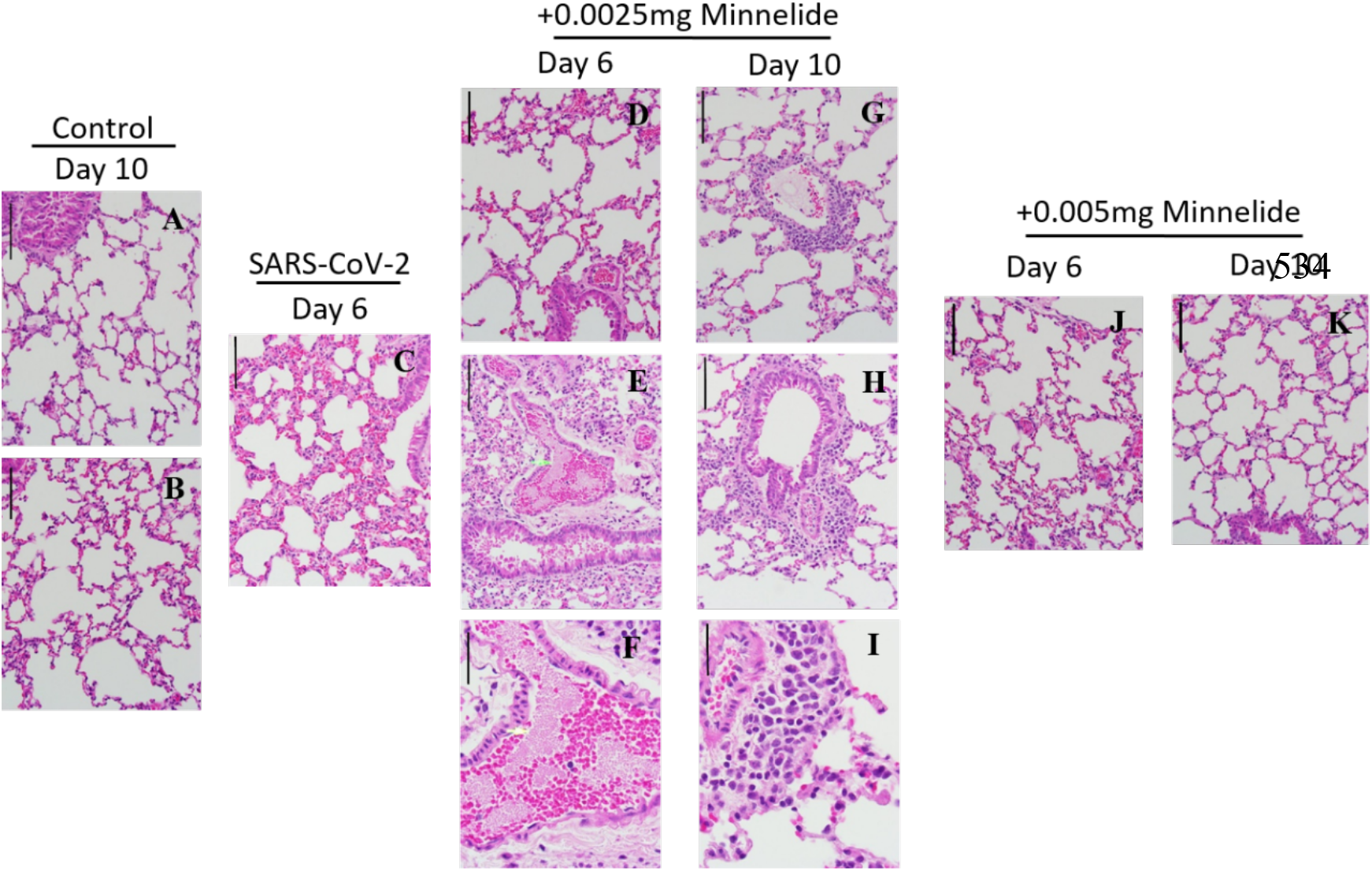
Histopathological Anaylsis of Mice after SARS-CoV-2 infection and Treatment with Minnelide. Histopathology of representative hematoxylin-and-eosin-stained, lung sections from Vehicle control mice at day 10 post-infection (A) High dose Minnelide (0.005mg) treated mice at day 10 post-infection (B) SARS-CoV-2 infected non-treated mice at day 6 post-infection (C) SARS-CoV-2 infected mice treated with low dose Minnelide (0.0025mg) at day 6 post-infection (D-F) and at day 10 post-infection (G-I) SARS-CoV-2 infected mice treated with high dose Minnelide (0.005mg) at day 6 post-infection (J) and day 10 post-infection (K). (A-E, G-H, and J-K) Scale bar 50μm and (F and I) scale bar 20μm.

## 3 Discussion

Beginning in December 2019, the COVID-19 pandemic rapidly created a global health emergency and killed millions of people worldwide. The clinical manifestation of this disease appears to be extremely variable, from asymptomatic to severe pneumonia leading to multi-organ failure requiring critical care. Recent studies indicate that a SARS-CoV-2 induced ARDS contributes to disease severity and mortality among patients. Although inflammation is a crucial host mechanism in preventing disease, in patients with COVID-19, hyperinflammation after SARS-CoV-2 infection causes a majority of the host tissue damage. Therefore, therapeutic approaches to control inflammation can be a way forward in limiting the damage caused by SARS-CoV-2 infection. Minnelide is a water soluble pro drug of a compound triptolide which was purified from a Chinese herb *Tripterygium wilfordii* Hook F (TWHF) [19]. For the past two thousand years, TWHF has been used in traditional Chinese medicine for the treatment of leprosy and rheumatoid arthritis. Further research revealed that triptolide is the bioactive component of TWHF which is responsible for its immunosuppressive anti-inflammatory properties. Unpublished work from our lab has shown reduced inflammation in a mouse model of chronic pancreatitis, a chronic inflammatory disease of pancreas, thereby making it a potential treatment for COVID-19. Our current study showed that Minnelide at a dose of 0.005mg/day in K18-hACE2-transgenic mice provide 100% survival benefit 10 days post-intranasal infection with SARS-CoV-2 (Figure 3). Further, we observed 100% mortality in nontreated mice post SARS-CoV-2 by day 6 post-infection (Figure 3).

Immunopathology of COVID-19 has revealed that immunity-mediated cytokine release syndrome is central to ARDS in patients with SARS-CoV-2 [27]. Initial results showed that patients with SARS-CoV-2 have higher level of proinflammatory cytokines in circulation such as IL-1α, IL-1β, IL-1 receptor antagonist protein (IL1RA), IL-2, IL-4, IL-6, granulocyte-macrophage colony-stimulating factor (GM-CSF), and IFN-gamma compared to healthy controls [4, 7]. In addition, patients with severe COVID-19 also have higher levels of IL-6, IL-8, and IL-1β in the BALF [8]. Furthermore, patients who have been admitted to Intensive Care Units (ICUs) have a higher level of proinflammatory cytokines than the non-ICU patients, which highlights the role of proinflammatory cytokines in severity of the disease. Our data shows that Minnelide treatment in K18-hACE2-transgenic mice infected with SARS-CoV-2 promotes survival by eliciting an anti-inflammatory response (Figure 4). Cytokine analysis of lungs revealed significant reduction in proinflammatory cytokine IL-6 after treatment with Minnelide (Figure 4).

D-dimer is the product of fibrinolytic degradation of fibrin, and elevated levels indicate the activation of coagulation and fibrinolysis systems in the body, which is extremely useful for the diagnosis of thrombotic diseases [28]. Healthy individuals have low levels of circulating D-dimer while chronic inflammatory conditions like rheumatoid arthritis, sickle cell disease, and asthma have elevated D-dimer level in the blood [29]. Increased D-dimer levels were also reported in patients with COVID-19 with even higher levels among COVID-19 patients who required hospitalization, as well as those aged older than 60 years [29]. Recent reports showed that respiratory deterioration is linked to thrombosis and high D-dimer level associated with poor prognosis and higher mortality in COVID-19 patients [30]. Fibrinogen is a soluble glycoprotein present in blood plasma which is enzymatically converted by thrombin to fibrin at the time of tissue injury, causing formation of fibrin based blood clot to stop bleeding. Recent studies shows that COVID-19 patients have significantly high fibrinogen levels in critically ill patients making them susceptible for the coagulopathy and prothrombic diathesis [31]. Therefore, monitoring levels of D-dimer and fibrinogen in COVID-19 patients helps in understanding the progressive severity of the disease with higher level predicting requirement of critical care and any experimental therapy against COVID-19. Our study shows that Minnelide significantly reduces D-dimer and fibrinogen level in lungs of SARS-CoV-2 infected mice (Figure 4). Furthermore, analysis of serum shows that Minnelide treatment causes a significant reduction in D-dimer level at day 6 post-infection in K18-hACE2-transgenic mice after SARS-CoV-2 infection when compared to the non-treated control (Figure 4).

Histologic evaluation of COVID-19 patients’ lung shows diffuse alveolar damage with hyaline membranes, fibrin deposits, edema multinucleated cells, and immune infiltration. Previous work by Yinda, et al. [18] has shown that K18-hACE2 mice develop edema-associated acute lung injury after SARS-CoV-2 infection which resembles the clinical features of COVID-19 patients. Analysis of hematoxylin and eosin-stained lung sections from K18-hACE2 mice infected with SARS-CoV-2 revealed that Minnelide treatment significantly improves lung histology on day 10 post-infection with all the mice showed normal lung tissue with thin alveolar septa with no inflammatory cells (Figure 5).

Minnelide is in clinical trials for the treatment of various cancers and doses used in present study are equivalent to human doses, which did not show any significant toxicity in a dose-escalation phase I clinical trial. In the current study, Minnelide, water soluble pro-drug of triptolide, shows significant improvement in survival in K18-hACE2 mouse model of SARS-CoV-2 infection. Further analysis reveals that Minnelide treatment induces anti-inflammatory response and showed significant improvement in lung histology. Therefore, present study highlights the potential of Minnelide in improving survival of hospitalized patients who have presented with ARDS after SARS-CoV-2 infection and require aggressive critical care.

## 4 Materials and Methods

### 4.1 Mice

The K18-hACE2 COVID-19 transgenic mouse model (B6.Cg-Tg (K18-ACE2)2Primm/J, Stock No. 034860, Jackson Laboratory, Bar Harbor, ME, USA) was used in the current study. These mice express the human angiotensin-converting enzyme 2 (hACE2) transmembrane protein under an epithelial cell promoter, rendering them susceptible to infection with human coronaviruses, such as SARS-CoV and CoV-2, that utilize hACE2 for attachment and entry into host cells. hACE2 male and female mice (6-8 weeks) were used for these studies according to NIH guidelines for housing and care in a biosafety level 3 animal laboratory. All procedures were approved by the Institutional Animal Care and Use Committee (protocol number 20-005) of Northern Arizona University.

### 4.2 Virus and Media

This study used coronavirus SARS-CoV-2 isolate USA-WA1/2020 (NR-52281; BEI Resources, Manassas, VA, USA). Viral propagation occurred in Vero E6 cells (ATCC CRL-1586) from the American Type Culture Collection (ATCC, Manassas, VA, USA) in Eagle’s minimum essential medium (EMEM) supplemented with 2% or 10% (for cell propagation) fetal bovine serum (FBS), 100 U/ml penicillin, 100 μg/ml streptomycin (Pen-Strep), 0.01 M HEPES, 1 mM sodium pyruvate, 1× nonessential amino acids solution (Thermo Fisher, USA), and 2 mM L-glutamine. Briefly, a t175 flask that was ~50% confluent was infected with SARS-CoV-2 virus, whereby 10 mL of fresh 2% FBS growth media containing 100μL of the BEI viral stock was added to VeroE6 cells and gentle rocked on a rocking platform for 1 h at room temperature. 15 mL of additional 2% FBS growth media was then added to the flask to bring the final volume to 25 mL. The flask was incubated for 7 days at 37°C in 5% CO_2_ and inspected for viral replication at day 7; cell death/rounding was used as a proxy for successful viral propagation and was observed by inverted microscope. The cells/virus were then subjected to three freeze/thaw cycles at −80°C to maximize the release of virus from cells. The contents were removed from the flask and placed in a 50mL conical tube that was centrifuged at 500 × *g* at 4°C for 15 min to separate the cellular debris from the supernatant containing the virus. 1 mL aliquots of the supernatant (viral stocks) were stored at −80°C for future use.

Concentration of the viral stocks were determined by plaque assay. 6-well plates (CLS3516; Millipore Sigma) were seeded with ~3.0 × 10^5^ cells/well and incubated for 48 to 72 h at 37°C in 5% CO_2_ until 80 to 90% confluency was reached. The viral stock was serially diluted in serum free Dulbecco’s MEM (DMEM; Millipore Sigma, USA). Serial dilutions were prepared as follows; 30μL of stock virus was added to 270μL DMEM to create a 10^−1^ stock, 30μL of this stock was then added to 270μL of DMEM to create a 10^−2^ stock. This continued to create 10^−3^, 10^−4^, and 10^−5^ viral stocks. Prior to infection, the growth medium was replaced with fresh medium containing 2% FBS and infected with 100μL of each SARS-CoV-2 dilution; dilutions 10^−1^ through 10^−5^ were tested in triplicate. The 6-well plates were briefly rocked on a rocking platform and set to incubate for 1 hour at 37°C in 5% CO_2_. Medium was then replaced with a 1× Dulbecco’s MEM (DMEM, Millipore Sigma, USA)/1.2% low-melting-point agarose (Bio-Rad, USA) overlay. This was allowed to solidify at room temperature for 15 min and incubated for 120 h at 37°C in a 5% CO_2_ atmosphere. Then, 2.0 ml of 4% paraformaldehyde was added to each overlay for 30 min, followed by staining with 1% crystal violet, removal of the overlay, and a triple rinse with phosphate-buffered saline (PBS). Uninfected cells were included as a control for all experiments and SARS-CoV-2 manipulations were conducted in a BSL-3 facility. PFUs were counted and averaged for each dilution. The concentration of undiluted viral stocks used for the animal experiments described herein was determined to be 1×10^5^ PFU/mL. Further dilution was completed in PBS to the infection concentration of 2000 PFU.

### 4.3 SARS-CoV-2 Infection and Treatments

K18-hACE2 mice were anesthetized with ketamine/xylazine (80/8 mg/kg) and intranasally inoculated with 2000 PFU SARS-CoV-2 suspended in 30 μl phosphate-buffered saline (PBS) and monitored for symptoms over a 10 day period. The day after infection, mice were treated with vehicle control (PBS), Minnelide low dose (0.0025mg/day) or high dose (0.005mg/day) via intraperitoneal injection. Disease presentations ranged from moderate (fur ruffling, lack of responses to stimuli, weight loss) to predetermined endpoint of 20% loss of pre-infection body weight of the animals. Mice were euthanized on pre-determined days post infection, or at end-point, with ketamine/xylazine (100/10 mg/kg) and cardiac puncture was performed to collect blood for cytokine analysis along with brain, lung, and heart excised using aseptic technique. Partial lung and heart were homogenized in 1 ml of sterile PBS for cytokine analysis. Brain, partial lung and heart were processed for histology and/or measurement of viral load using RT-qPCR. For animals that reached humane euthanasia endpoints at any time, or those surviving the duration of the experiments, the brain, heart, lung, and serum were obtained at necropsy and processed for histology, ELISA, and measurement of viral load using RT-qPCR, respectively.

### 4.4 Reverse transcriptase quantitative real-time PCR (RT-qPCR) of viral genomic RNA

The left lobe and partial brain from each mouse was weighed (0.05-100mg) and placed in a prefilled bead tube containing 1.00mm glass beads (Beadbug, Z763756) and 600μl of lysis buffer with 1% BME from the Purelink RNA mini kit (Invitrogen, USA). The tissue was homogenized for 1 minute disruption at 5 m/s and 1 min cooling in ice, repeated twice. Tubes were spun at 2000g for 5 minutes at 4°C. RNA was isolated using kit directions including incorporation of homogenizer tubes (Invitrogen, 12183026) to remove remaining animal tissue.

One step RT-qPCR was conducted to detect the viral RNA load in each sample using Reliance One-Step Multiplex Supermix (BioRad, USA) containing the Reliance Reverse Transcriptase enzyme. Primers CoV2-S_19F (5’ -GCTGAACATGTCAACAACTC-3’) and CoV2-S_143R (5’ - GCAATGATGGATTGACTAGC-3’) were designed to target a 125bp region of the SARS-CoV-2 spike protein. Probe CoV2-S_93FP (5’ – ACTAATTCTCCTCGGCGGGC-3’) was fluorescently labeled with FAM dye for RT-qPCR detection using the QuantStudio 12K Flex Real-Time PCR System, as previously described (32). Each qPCR was prepared as a 20μL reaction containing 12.5μL molecular grade H_2_O, 5μL 4X Reliance Supermix (1x final concentration), 0.2μL forward primer, 0.2μL reverse primer, and 0.1μL probe, with 2μL template RNA. During the RT-qPCR, reverse transcription occurred at 50°C for 10 minutes followed by a hot start step at 95°C for 10 minutes. This was followed by denaturation at 95°C for 10 seconds and annealing at 60°C for 30 seconds which were repeated for a total of 40 cycles. Viral copy number was extrapolated using synthetically generated internal qPCR standards as previously described (32).

### 4.5 Cytokine Analysis

Serum was collected in BD Vacutainer™ tubes (ThermoFisher, USA). Samples were processed by centrifuging 3500 RPM for 5-10 minutes, supernatant collected, and protease inhibitor added (Halt™ cocktail, ThermoFisher, USA). Heart and lung were excised and homogenized in 1mL of ice-cold sterile PBS, an antiprotease buffer solution containing PBS, protease inhibitors (inhibiting cysteine, serine, and other metalloproteinases), and 0.05% Triton X-100 was added to the homogenate and then clarified by centrifugation 3500 RPM for 10 minutes. Supernatants were assayed for the presence of IL-6, which was quantified using an Invitrogen ELISA kit (ThermoFisher, USA); and DDimer and Fibrinogen were quantified using Biomatik ELISA kits (Biomatik Corporation, Ontario, Canada).

### 4.6 Histology

Tissues were fixed in histology cassettes suspended in 10% formalin (Sigma Aldrich) for 48 hours. Histology cassettes of fixed tissues were then fixed using the routine overnight non-fat cycle on the Excelsior AS (Thermo Scientific) and embedded in paraffin wax (Histoplast LP, Fisher Scientific). Tissues were sectioned in 5μm thin sections onto charged slides and heat fixed overnight at 45°C. Sections were then processed for H&E staining and submitted to pathologist for analysis.

### 4.7 Statistical Analysis

Differences in data were tested for significance using either GraphPad Prism for Windows (GraphPad, USA). For details of the statistical test applied, refer to the figure legends. P < 0.05 were considered significant.

## 5 Acknowledgements

Funding provided by Minneamrita Therapeutics LLC to NAU.

